# Measurement of leaf day respiration using a new isotopic disequilibrium method compared with the Laisk method

**DOI:** 10.1101/201038

**Authors:** Xiao Ying Gong, Guillaume Tcherkez, Johannes Wenig, Rudi Schäufele, Hans Schnyder

## Abstract

- Quantification of leaf respiration is of great importance for the understanding of plant physiology and ecosystem biogeochemical processes. Leaf respiration continues in light (*R*_L_) but supposedly at a lower rate compared to the dark (*R*_D_). Yet, there is no method for direct measurement of *R*_L_ and most available methods require unphysiological measurement conditions.
- A method based on isotopic disequilibrium quantified *R*_L_ (*R*_L 13C_) and mesophyll conductance of young and old fully-expanded leaves of six species compared *R*_L 13C_ to *R*_L_ values determined by the Laisk method (*R*_L Laisk_).
- *R*_L 13C_ and *R*_L Laisk_ were consistently lower than *R*_D_. Leaf ageing negatively affected photosynthetic performance, but had no significant effect on *R*_L_ or *R*_L_/*R*_D_ as determined by both methods. *R*_L Laisk_ and *R*_L 13C_ were measured successively on the same leaves and correlated positively (*r*^2^=0.38), but average *R*_L Laisk_ was 28% lower than *R*_L13C_. Using *A*/*C*_c_ curves instead of *A*/*C*_i_ curves, a higher photocompensation point Γ* (by 5 μmol mol^-1^) was found but the correction had no influence on *R*_L Laisk_ estimates.
- The results suggest that the Laisk method underestimated *R*_L_. The isotopic disequilibrium method is useful for assessing responses of *R*_L_ to irradiance and CO_2_, improving our mechanistic understanding of *R*_L_.

## INTRODUCTION

Foliar respiration is a major component of the global carbon cycle, releasing more than three times the amount of CO_2_ liberated by anthropogenic emission each year (Le Quere *et al.*, 2009; Beer *et al.*, 2010), if it is assumed that foliar, i.e. plant leaf respiration constitutes 50-80% of plant respiration globally (Atkin *et al.*, 2007; Lehmeier *et al.*, 2010). Thus knowledge of the drivers and controls of leaf respiration is essential for understanding plant physiology and the global carbon budget, and that knowledge is required for improving the representation of leaf respiration in climate-vegetation models (Atkin *et al.*, 2007; Heskel *et al.*, 2013). The fact that leaf respiration rate is lower in light (*R*_L_, also termed day respiration) compared to the dark (*R*_*D*_) – when normalized to the same temperature – has long been recognized and demonstrated in leaf- (Brooks & Farquhar, 1985; Atkin *et al.*, 2000; Gong *et al.*, 2015), stand- (Schnyder *et al.*, 2003; Gong *et al.*, 2017a), and ecosystem-scale (Wehr *et al.*, 2016) studies. The inhibition of respiration by light is underpinned by the light-induced down-regulation of the activity of several enzymes of respiratory metabolism (Tcherkez *et al.*, 2005; Tcherkez *et al.*, 2012a). Yet, the quantification of *R*_*L*_ is technically challenging and the mechanism controlling its variation is uncertain.

In practice, *R*_L_ cannot be directly measured using conventional gas exchange measurements because *R*_L_ is masked by other concurrent, major fluxes: photosynthetic CO_2_ uptake and photorespiratory CO_2_ release. Net CO_2_ assimilation rate can be expressed as: *A* = *V*_c_ - 0.5 *V*_o_ - *R*_L_, where *V*_*c*_ is the rate of carboxylation and *V*_o_ is that of oxygenation, and 0.5 *V*_o_ is the rate of photorespiration (*F*). *A* can be further expressed as:

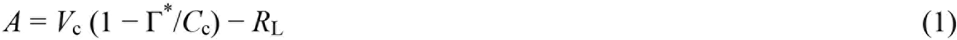

where Γ* is the CO_2_ compensation point in the absence of day respiration and *C*_c_ is the chloroplastic CO_2_ mole fraction. At a *C*_*c*_ that equals to Γ*, *A* is equal to –*R*_L_. Based on Eqn 1, *R*_L_ can be estimated from the common intersection of curves of net CO_2_ assimilation rate (*A*) vs. intercellular CO_2_ concentration (*C*_*i*_) measured under low CO_2_ and sub-saturating levels of irradiances (defined as *R*_L Laisk_ here), as described in Laisk (1977) and further extended by Brooks & Farquhar (1985). This is based on the notion that at *R*_L_ is insensitive to light intensity. The Laisk method uses *A*/*C*_i_ curves instead of *A*/*C*_c_ curves to determine the common intersection, and thus gives the apparent, *C*_i_-based CO_2_ compensation point 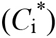 and *R*_L_ at 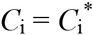. Although it has been widely used as a standard method for determining *R*_L_ and Γ* (in assuming 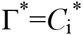) (Farquhar *et al.*, 1980; von Caemmerer, 2000; Walker & Ort, 2015), uncertainties and limitations of the Laisk method have been intensively discussed. First, ignoring the influence of mesophyll conductance (*g*_m_) might lead to errors in estimates of *R*_L_ and Γ*, as 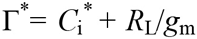 (Brooks & Farquhar, 1985; von Caemmerer *et al.*, 1994; Walker & Ort, 2015). Second, the measurement must be performed at very low CO_2_ that generally contrast with growth conditions (Villar *et al.*, 1994; Yin *et al.*, 2011). Experimental evidence has indicated a CO_2_ effect on respiration rate in light (Gong *et al.*, 2017a) and on the abundance of transcripts encoding enzymes of the respiratory pathway in both long-term (Leakey *et al.*, 2009) and short-term (Li *et al.*, 2013) treatments. These observations raise the concern that *R*_L_ measured by the Laisk method might differ from actual *R*_L_ under growth conditions. Similarly, other methods, such as the Kok method (Kok, 1948) and a method based on chlorophyll fluorescence (Yin *et al.*, 2011), generally must be performed at low CO_2_ levels or low irradiance levels and require manipulation of CO_2_ assimilation rate (for a review see Yin *et al.*, 2011). Furthermore, during both Kok and Laisk measurements, variations in *C*_c_ are critical but have not been accounted for, potentially leading to errors in *R*_L_ estimates (Farquhar & Busch, 2017; Tcherkez *et al.*, 2017a b).

Techniques that allow measuring *R*_L_ without requiring modifications of environmental conditions such as CO_2_ mole fraction or irradiance typically use carbon isotopes. The principle of deconvoluting CO_2_ flux components by artificially created isotopic disequilibrium (i.e. labelling) has been widely explored for i.e. photorespiration (Ludwig & Canvin, 1971) or stand- (Schnyder *et al.*, 2003; Gong *et al.*, 2017a) or ecosystem-scale (Ostler *et al.*, 2016) autotrophic respiration. This type of labelling method exploits the fact that CO_2_ flux components have distinct dynamics of tracer incorporation during the labelling. Abrupt changes to a ^13^CO_2_ atmosphere were used to monitor the liberation of ^12^CO_2_ by respiration in the first minutes following the isotopic changeover (Loreto *et al.*, 2001; Pinelli & Loreto, 2003). However, when using pure ^13^CO_2_ this technique is relatively costly and requires a ^13^C-sensitive infrared gas analyzer. Gong *et al.* (2015) described a leaf-level isotopic disequilibrium method to quantify *R*_L_ using CO_2_ sources of natural ^13^C abundance, which is based on concurrent measurements of photosynthetic gas exchange and ^13^C/^12^C isotope composition (denoted as δ, definition see methods) of CO_2_ fluxes, i.e. online ^13^C discrimination by net photosynthesis (online Δ). In other words, the δ-value of gross fixed CO_2_ (associated with the flux *V*_c_) responds instantaneously at the onset of labelling (i.e. abrupt change of δ of CO_2_ fed to leaf), with the δ-value of the photorespired CO_2_ (flux 0.5 *V*_o_) following with only a short delay (half-life in the order of a few minutes (Ludwig & Canvin, 1971)). By contrast, the δ-value of respired CO_2_ responds rather slowly (half-life in the order of one to a few days (Schnyder *et al.*, 2003; Lehmeier *et al.*, 2008; Tcherkez *et al.*, 2012b; Gong *et al.*, 2017a). This approach requires two sets of online Δ measurements on similar leaves (or the very same leaves, as in this study), so as to examine the isotopic mass balance at the photosynthetic steady-state (Gong *et al.*, 2015). This method has the following advantages: (i) *R*_L_ measurements can be done at any setting of environmental parameters, e.g. identical to growth conditions; (ii) it measures *R*_L_ at the photosynthetic steady-state without manipulation of the photosynthesis rate; (iii) it simultaneously provides a reliable measurement on mesophyll conductance (*g*_m_), another important parameter. As it relies on the measurements of δ-values and CO_2_ exchange rates, diffusive leaks across the gasket of leaf cuvette must be minimized (Gong *et al.*, 2017b) or accounted for (Gong *et al.*, 2015).

Here, we use the isotopic disequilibrium method (presented by Gong *et al.* 2015) to measure *R*_L_ (*R*_L 13C_) on single leaves, and compare the results with the Laisk method (*R*_L Laisk_) applied to the very same leaves. Thus, our objectives were to answer the following questions. (i) Does the isotopic disequilibrium method also show an inhibition of leaf respiration by light? (ii) Do *R*_L_ estimates from isotopic disequilibrium agree with those from the Laisk method for different species and leaves of different age effects? Or (iii) is there any consistent offset in *R*_L_ estimates obtained with the two methods, and if yes, is the offset correlated to leaf age or simply due to assumptions on internal/mesophyll conductance? To this end, ^13^CO_2_/^12^CO_2_ exchange of leaves from plants grown with ambient CO_2_ with a δ^13^C of CO_2_ (δ^12^C_Co2_) near −10‰ was measured sequentially in the presence of CO_2_ with a δ^13^C_CO2_ of −31.2‰ and −6.3‰, and *R*_L_ of leaves was solved using isotopic mass balance equations. These measurements were immediately followed by determinations of *R*_L Laisk_. The comparison of *R*_L 13C_ and *R*_L Laisk_ was performed on both young and old mature leaves of two grass and four legume species. Villar *et al.* (1995) have reported that ageing of leaves of an evergreen shrub led to a reduction of *R*_L Laisk_/*R*_D_ from 0.5 to 0.2. This is the reason why we included young and old leaves, since it might increase the variation range of *R*_L_ and thus enhance the method comparison. In addition, we estimated *g*_m_ of every leaf, so that *A*/*C*_i_ curves could be converted to *A*/*C*_c_ curves to estimate Γ* and *R*_L Laisk_ based on the common intersection of *A*/*C*_c_ curves.

## MATERIALS AND METHODS

### Plant material and growth conditions

Six herbaceous plant species were used, namely barley (*Hordeum vulgare*), wheat (*Triticum aestivum*), castor bean (*Ricinus communis*), French bean (*Phaseolus vulgaris*), soybean (*Glycine max*) and broad bean (*Vicia faba*). Plants were grown from seed in plastic pots filled with quartz sand, placed in a growth chamber (PGR15, Conviron, Winnipeg, Canada) and supplied with a modified Hoagland nutrient solution with 7.5 mM nitrate (cf. Gong *et al.*, 2017b) every two to three days. Environmental conditions during plant growth were: a photosynthetic photon flux density (PPFD) of 700 μmol m^-2^ s^-1^ during the 12 h-long photoperiod per day, ambient CO_2_ concentration ([CO_2_]) of about 400 μmol mol^-1^, air temperature of 22 °C during photo- and dark- periods, relative humidity of 50% during photoperiod and 60% during dark period. The density of plants in the growth chamber was rather low, thus leaves were not shaded. Young leaves, defined as the youngest fully expanded leaves, were measured when plants reached a stage of having 3-4 mature leaves per branch/tiller. Old leaves, defined as two age categories older than the measured young leaves, were measured about 10 days later. At that time plants had 5-7 mature leaves per branch/tiller. For dicots, the fully expanded terminal leaflets were measured. Young leaves were measured for all species, while old leaves of *G. max* and *V. faba* were not measured.

### ^13^CO_2_/^12^CO_2_ gas exchange facilities

^13^CO_2_/^12^CO_2_ gas exchange and labelling were performed using the protocols and facilities described in Gong *et al.* (2015) with modifications and advancements as follows. The approaches in Gong *et al.* (2015) provided a mean leak coefficient and a *R*_L_/*A* for a group of similar leaves (same species and age, treated as replicates). In this study, leak coefficients were measured for each leaf and used for the correction of its gas exchange data, using the equations in Gong *et al.*, (2015). To quantify *R*_L_, the two components of *A* must be separated, as *A* = *P* – *R*_L_, where *P* is the apparent photosynthesis rate (*P* = *V*_c_ – *F*). Briefly, we switch the CO_2_ source supplied to leaf photosynthesis to create isotopic disequilibrium between *P* and *R*_L_, namely, *P* will be immediately labelled while *R*_L_ is fed by substrate formed during plant growth (old carbon) (Gong *et al.*, 2015), thus *R*_L_ can be solved by isotopic mass balance (see below).

The leaf-level ^13^CO_2_/^12^CO_2_ gas exchange and labelling system included a portable CO_2_ exchange system (LI-6400, LI-COR Inc., Lincoln, USA) housed in a gas exchange mesocosm (chamber 1, cf. Gong *et al.*, 2015; Gong *et al.*, 2017b), and another gas exchange mesocosm (chamber 2) for the purpose of providing labelling CO_2_. The air supply to both mesocosms and the LI-6400 was mixed from CO_2_-free, dry air (with 21% O_2_) and CO_2_ of known δ^13^C_CO2_ (cf.(Schnyder *et al.*, 2003), with δ^13^C denoting the ^13^C composition of a sample defined as the relative deviation of its ^13^C/^12^C ratio (ℜ_sample_) to that of the international VPDB standard (ℜ_VPDB_): δ^13^C = ℜ_sample_/ℜ_VPDB_ − 1. [CO_2_] inside chamber 1 was monitored with an infrared gas analyzer (LI-6262, LI-COR Inc., Lincoln, USA). During leaf gas exchange measurements, the plants to be measured and the sensor head of the LI-6400 were placed inside the chamber 1. Using this setup,we separately controlled the CO_2_ concentration and δC_co2_ in the leaf cuvette and growth chambers. The growth chamber and leaf cuvette systems were coupled to a continuous-flow stable isotope ratio mass spectrometer (IRMS; Delta^plus^ Advantage equipped with GasBench II, ThermoFinnigan, Bremen, Germany) for ^13^C analysis of the sample air. The whole-system precision of repeated measurements on δ^13^C was 0.09‰ (SD, *n*=50). For further details of the method see Gong *et al.* (2015) and Gong *et al.* (2017b).

### Determinations of *K*_C02_, *R*_D_

Measurements of each leaf started with the determination of the cuvette leak coefficient for CO_2_ (*K*_CO2_) with the leaf present in the cuvette during these measurements (Gong *et al.*, 2015). Each leaf was held in the leaf cuvette of the LI-6400 for more than 20 min in the dark, at a constant [CO_2_] of 488 ± 9 (SD) μmol mol^-1^ in the leaf cuvette (*C*_out_) and 400 μmol mol^-1^ in the chamber 1 (*C*_M_) that housed the LI-6400 measurement head (detailed measurement conditions are shown in Table S1). When gas exchange had reached a constant rate, gas exchange parameters, including [CO_2_] and the δ^13^C of the incoming (*C*_in_ and δ_in_) and outgoing cuvette air (*C*_out_ and δ_out_) were measured with the LI-6400 and the online IRMS. Thereafter, *C*_M_ was reduced to about 200 μmol mol^-1^ and the same gas exchange parameters were measured at steady-state. Since manipulating *C*_M_ should only affect the diffusive leak between the chamber 1 housing the leaf gas exchange equipment and the internal space of the leaf cuvette but not *R*_D_, *K*_CO2_ was determined as the slope of the observed net CO_2_ exchange rate in the dark (*N*_D_) and (*C*_M_ − *C*_out_)/*s* relationship as:

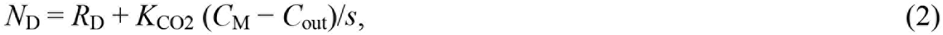

where *s* is the leaf area (Gong et al. 2015). Knowing the *K*_CO2_ of each intact leaf, CO_2_ exchange data were corrected as shown in Gong *et al.* (2015) and *R*_D_ determined. Since leak coefficients for ^12^CO_2_ and ^13^CO_2_ were virtually the same (Gong *et al.*, 2015), *K*_CO2_ was used to correct both ^12^CO_2_ and ^13^CO_2_ flux data. Before all calculations, data of δ and rates of CO_2_ fluxes were corrected for leak artefact using *K*_CO2_ of individual leaves and equations in Gong *et al.* (2015).δ^13^C of *R*_D_ was calculated as:

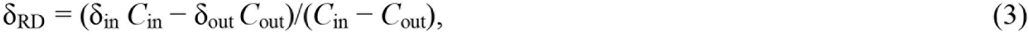

with δ_in_ and δ_out_ are δ measured at inlet and outlet air stream, respectively.

### ^13^C labelling

After measurements of *K*_CO2_, *R*_D_ and δ_RD_, the light source of the LI-6400 was switched on (PPFD 700 μmol m^-2^ s^-1^) to measure the online Δ using CO_2_ sources with different δ^13^C of CO_2_ (−6.3‰ and −31.2‰). The ^12^C/^13^C discrimination associated with net photosynthesis, Δ_A_, was calculated according to Evans *et al.* (1986): Δ_A_ = ξ(δ_out_ − δ_in_)/(1+ δ_out_ − ξ(δ_out_ − δ_in_)), where ξ = *C*_in_/(*C*_in_ − *C*_out_). Here, ξ was below 15 during Δ_A_ measurements. Measurements of Δ_A_ were done in the photosynthetic steady-state: after about 30 min of stabilization in the conditions similar to that of plant growth average *C*_out_ of 394 ± 34 (SD) μmol mol^-1^, average relative humidity of 76 ± 10 %, block temperature of 22 °C (mean leaf temperature was 23.3 ±0.2 °C, Table S1). Online Δ was firstly measured using the depleted CO_2_ source (−31.2‰), then measured with the enriched CO_2_ source (−6.3‰) on each leaf. Chamber 2 was used to mix the labelling air containing the enriched CO_2_ with the targeted [CO_2_]. When labelling start, well mixed air in Chamber 2 was supplied to the inlet of LI-6400 with a peristaltic pump. Using this setup, the labelling air can completely flush out the air in the LI-6400 system within 8 min. The second online Δ was measured within 15 min after the start of labelling (i.e. switching of CO_2_ sources), and all photosynthetic gas exchange rates are not influenced by labelling, as only δ^13^C of CO_2_ fed to leaf was changed (Gong *et al.*, 2015).

### Calculations of *R*_L_

Substituting the relationship giving the photosynthetic assimilation in the absence of day respiration *P* (= *V*_c_ − *F*) into equation (1) gives:

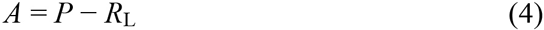

Applying isotopic mass-balance to equation (4) gives:

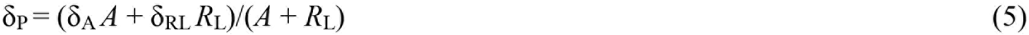

where δ_P_, δ_A_, δ_RL_ are the δ^13^C of *P*, *A* and *R*_L_, respectively. With the two sets of online Δ measurements we have:

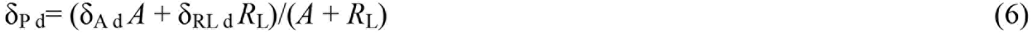

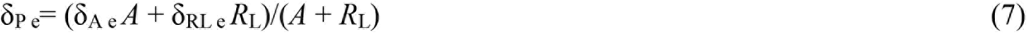

where subscripts “d” and “e” indicates parameters measured with the ^13^C-depleted and ^13^C-enriched CO_2_ sources, respectively. Since ^13^C discrimination in *P* (Δ_P_), is independent of the δ^13^C of the CO_2_ source (Farquhar et al., 1989):

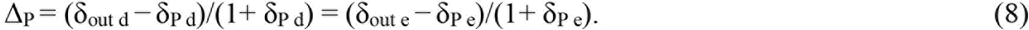

Combining the rearranged Eqn 6-8 we have:

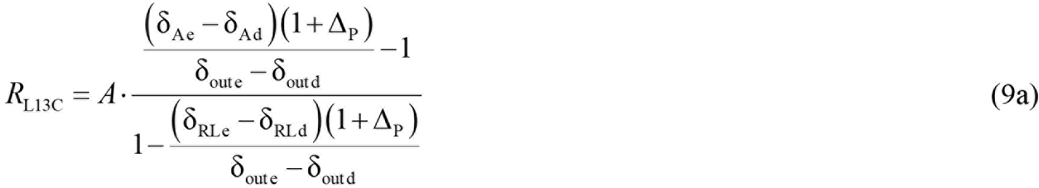

Equation (9a) includes the isotope composition of day-respired CO_2_ (both under a ^13^C-enriched and ^13^C-depleted atmosphere) in the denominator. Under the assumption that day respiration reacts very slowly to photosynthetic input (see *Introduction*), δ_RL d_ = δ_RL e_ and equation (9a) rearranges to:

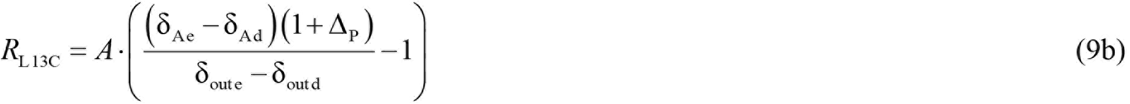

In practice, the approximation δ_RL d_ = δ_RL e_ is not critical: if some C atoms photosynthetically fixed under the ^13^C-depleted atmosphere were channelled to respiratory metabolism and liberated as CO_2_ under the ^13^C-enriched atmosphere, this would lead to a change of a few per mils only in the denominator and the change in *R*_L 13C_ would be very small. In fact, during the first measurement phase (≈ 20 min) under the ^13^C-depleted atmosphere, we expect at most 10% turnover in leaf respiratory pools (measured by Nogues *et al.* (2004) for dark respiration) meaning a maximal putative change in δ_RL_ of about 0.6‰ (Table S2, the denominator in equation 9a would thus be equal to 0.975 instead of 1).

In Gong *et al.* (2015), the approximation that 1+Δ_P_=1 was used. Here, we applied a different approximation that Δ_P_ = Δ_A e_, which was shown to be an acceptable approximationwhen the enriched CO_2_ source (−6.3‰) was close to that of the growth environment (−10‰) (Gong *et al.*, 2015).

Thus, *R*_L_ was calculated as follows:

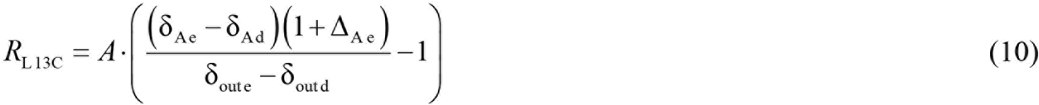

The δ-value of net assimilated CO_2_ was calculated as: δ_A_ = (δ_in_ *C*_in_ − δ_out_ *C*_out_)/(*C*_in_ − *C*_out_). *R*_L 13C_/*R*_D_ was calculated with a (small) correction accounting for the temperature difference between light and dark, using a Q_10_ of 2 (see Gong *et al.*, 2015).

### Calculation of mesophyll conductance

Mesophyll conductance (*g*_m_) is defined as *g*_m_ = *A*/(*C*_i_ − *C*_c_) (cf. Evans *et al.*, 1986), where *C*_c_ is the CO_2_ mole fraction at the site of carboxylation in the chloroplast. Estimation of *C*_c_ was based on the photosynthetic ^12^C/^13^C discrimination model of Farquhar *et al.* (1989) (cf. Gong *et al.*, 2015). In fact, a modified equation of ^13^C/^12^C discrimination that includes both mesophyll resistance and ternary effects (Farquhar & Cernusak, 2012) is:

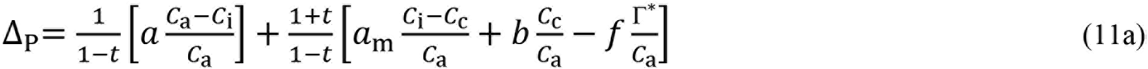

while the simplified equation that excludes mesophyll resistance (or assumes infinite *g*_m_) can be written as:

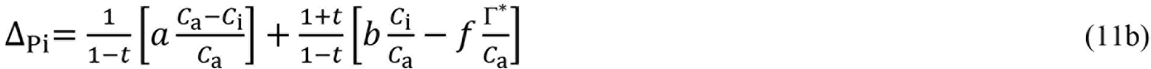

Therefore, the subtraction (11a) - (11b) gives:

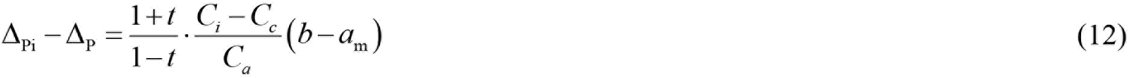

where *a* = 4.4‰, *b* = 28.9‰, *a_m_* combines dissolution and diffusion in the liquid phase so that *a*_m_ = 1.8‰ (Evans *et al.*, 1986) and *f* = 11‰ (Ghashghaie *et al.*, 2003; Lanigan *et al.*, 2008). Γ*, was approximated to be equal to 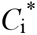 measured by the Laisk method (see below). *t* represents the ternary correction factor (Farquhar & Cernusak, 2012):

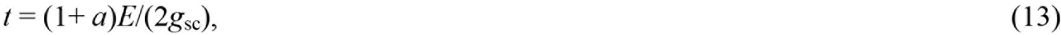

where *E* is the transpiration rate and *g*_sc_ is the stomatal conductance to CO_2_. Here, we ignored the boundary layer resistance because air was well mixed in the leaf cuvette of the LI-6400 (Kromdijk *et al.*, 2010). Δ_p_ can be calculated using Eqn 6-8, assuming δ_RL_ = δ_RD_ (Gong *et al.* 2015). Each leaf had two measurements of Δ_p_ using the two CO_2_ sources (Δ_p e_ and Δ_p d_, and theoretically they should be very similar and can be treated as technical replicates, see also Fig. 3), thus the mean of Δ_p e_ and Δ_p d_ was used to calculate *C*_c_ using Eqn 12.

It should be noted that equation (11a) simply represents the model of photosynthetic fractionation where the term associated with day respiration has been omitted. That is, the full model following Farquhar *et al.* (1989) notations is:

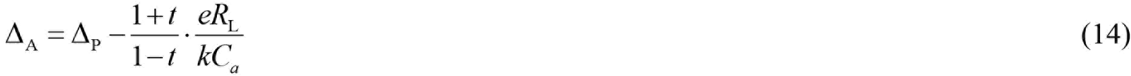

where *e* is the isotope fractionation by day respiration, with respect to net fixed photosynthates which are assumed to represent the respiratory substrates. However, considering that respiratory substrate pool turn-over is slow and mostly disconnected from photosynthesis at time scales less than the duration of the measurements (30-45min), day respiration is fed by a distinct carbon source and thus equation (14) has to be changed to (Tcherkez *et al.*, 2011):

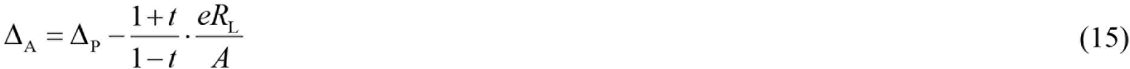

In Eqn 15, *e* is still expressed relative to net fixed CO_2_ (i.e. *e* = (δ_A_ − δ_RL_)/(δ_RL_ + 1)). Under our conditions, *t* is very small (< 0.1‰), thus Eqn 15 can be rearranged as

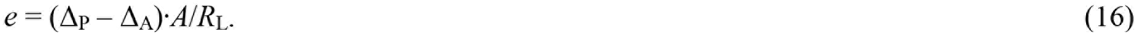

### Measurement of *R*_L_ and 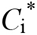 using the Laisk method

After online Δ measurements, each single leaf was measured for *R*_L_ using the Laisk method (Laisk, 1977; Brooks & Farquhar, 1985) with the LI-6400 open system. Briefly, *A*/*C*_i_ curves were obtained at three levels of PPFD, 50-70, 100-150, and 250 μmol m^-2^ s^-1^, and *C*_out_ was decreased from 110 to 50 μmol mol^-1^ step-wise at each PPFD. Average relative humidity was 77±9% and block temperature 22 °C (meaning that leaf temperature was 22.4±0.2 °C, Table S1). Again, the observed *A* and *C*_i_ values were firstly corrected for leak artefacts. The coordinates of the common intersection of *A*/*C*_i_ curves provided the estimates of *R*_L Laisk_ and 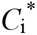 (Fig. S4). We also tested the slope-intersection regression approach suggested by Walker & Ort (2015), a modified Laisk method, but it yielded very similar results (data not shown) as the common intersection approach in the original Laisk method.

Using Laisk measurements, we estimated *R*_L Laisk CC_ and Γ* from the *A*/*C*_c_ curves (cf. Fig. S4). For this purpose, we established the relationship between *g*_sc_ and *g*_m_ across the measured leaves, and *g*_sc_/*g*_m_ was plotted against *A* or *C*_out_ to check whether the *g*_sc_ - to - *g*_m_ ratio was independent of photosynthesis rate or CO_2_ mole fraction. Using the *g*_sc_/*g*_m_ relationship, *g*_m_ along *A*/*C*_i_ curves was estimated from measured *g*_sc_, and thus *A*/*C*_i_ curves could be converted into *A*/*C*_c_ curves.

## RESULTS

### *R*_L_ *across species and leaf age*

As expected, both *R*_L Laisk_ and *R*_L 13C_ were consistently lower than *R*_D_, demonstrating that the labeling technique generally also shows an inhibition of leaf respiration in the light compared to the dark (Table 1). Further, leaf age had no effect on *R*_L Laisk_ or *R*_L 13C_ (Table 1). Also, both methods showed similar species effects: *H. vulgare* and *P. vulgaris* had higher *R*_L Laisk_ and *R*_L 13C_ than the other species; *T. aestivum* had the lowest *R*_L Laisk_ and *R*_L 13C_ of young leaves and *R. communis* the lowest *R*_L Laisk_ and *R*_L 13C_ of old leaves. Pooling over all *R*_L 13C_ and R_L Laisk_ paidata, a significant positive correlation was found (*r*^2^ =0.38, p<0.001, Fig. 1). Importantly, however, *R*_L Laisk_ was systematically smaller than *R*_L 13C_ by 28% (averaged over all leaves), and this effect was similar for the different species and age classes (Fig. 1). As a result, the ratio of respiration in light to that in darkness at the same temperature (*R*_L_/*R*_D_) was higher for the isotopic disequilibrium method than the Laisk method: *R*_L 13C_/*R*_D_ ranged between 0.6 and 1.3 with a mean of 0.9, and *R*_L Laisk_/*R*_D_ ranged between 0.4 and 0.9 with a mean of 0.7 (Fig. 2). Both measurements showed a tendency of increasing *R*_L_/*R*_D_ with leaf ageing; however, a significant age effect on *R*_L 13C_/*R*_D_ was detected in *P. vulgaris* while a clear age effect on *R*_L Laisk_/*R*_D_ was found in *P. vulgaris, T. aestivum* and *R. communis* (Fig. 2). *R*_D_ was not significantly different between age classes, but differed between species, with *T. aestivum* having the smallest *R*_D_ value of all species.

**Table 1.** Gas exchange parameters of young and old leaves measured under the same environmental conditions as during growth. Gas exchange parameters include: respiration rate in light measured using the isotopic disequilibrium method (*R*_L 13C_, μmol m^-2^ s^-1^) or the Laisk method (*R*_L Laisk_, μmol m^-2^ s^-1^),respiration rate in the dark (*R*_D_, μmol m^-2^ s^-1^),leak coefficient (*K*_CO2_, μmol s^-1^), net assimilation rate (*A*, μmol m^-2^ s^-1^), transpiration rate (*E*, mmol m^-2^ s^-1^), stomatal conductance to water vapour (*g*_sw_, mol m^-2^ s^-1^), mesophyll conductance (*g*_m_, mol m^-2^ s^-1^), apparent chloroplastic CO_2_ photocompensation point (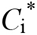 μmol mol^-1^), intrinsic water use efficiency (WUE_i_=*A*/*g*_sw_, μmol mol^-1^), ratio of internal to atmospheric CO_2_ concentration (*C*_i_/*C*_a_), ratio of chloroplastic to atmospheric CO_2_ concentration (*C*_c_/*C*_a_). Data are shown as mean±SE (*n*=4), significant treatment effects (age, species and their interaction, *a*×*s*) were marked with * when 0.01<*p*<0.05 or with ** when *p*<0.01.

|  | *H. vulgare* |  | *T. aestivum* |  | *P. vulgaris* |  | *R. communis* |  | *Significance* |  |  |
| --- | --- | --- | --- | --- | --- | --- | --- | --- | --- | --- | --- |
| | Young | Old | Young | Old | Young | Old | Young | Old | *age* | *species* | *a*×*s* |
| $R_{L\ 13C}$ | 1.8±0.1 | 1.7±0.3 | 1.0±0.1 | 1.4±0.2 | 1.5±0.1 | 2.2±0.2 | 1.5±0.1 | 1.0±0.1 | 0.45 | ** | * |
| $R_{L\ \text{Laisk}}$ | 1.1±0.1 | 1.4±0.2 | 0.7±0.1 | 0.9±0.1 | 1.3±0.1 | 1.7±0.3 | 0.8±0.2 | 0.7±0.1 | 0.25 | ** | 0.31 |
| $R_D$ | 1.9±0.2 | 1.8±0.1 | 1.0±0.2 | 1.0±0.1 | 2.3±0.1 | 2.2±0.2 | 2.0±0.4 | 1.1±0.1 | 0.07 | ** | 0.08 |
| $K_{\text{CO}_2}$ | 0.19±0.06 | 0.27±0.07 | 0.11±0.02 | 0.15±0.01 | 0.24±0.07 | 0.21±0.05 | 0.12±0.04 | 0.18±0.07 | 0.32 | 0.14 | 0.75 |
| $A$ | 14.1±0.5 | 7.5±1.7 | 15.4± 0.6 | 8.9±1.0 | 7.3±0.9 | 8.3±0.6 | 9.0±1.9 | 7.2±1.8 | ** | ** | ** |
| $E$ | 1.5±0.1 | 1.1±0.1 | 1.4±0.1 | 1.1±0.2 | 1.0±0.1 | 1.0±0.1 | 0.9±0.2 | 0.7±0.2 | 0.06 | ** | 0.71 |
| $g_{\text{sw}}$ | 0.26±0.03 | 0.11±0.02 | 0.37±0.06 | 0.11±0.03 | 0.21±0.08 | 0.12±0.02 | 0.16±0.09 | 0.08±0.03 | ** | 0.21 | 0.36 |
| $g_m$ | 0.25±0.02 | 0.08±0.03 | 0.32±0.04 | 0.12±0.02 | 0.09±0.02 | 0.10±0.01 | 0.12±0.03 | 0.08±0.03 | ** | ** | ** |
| $C_i^*$ | 41.8±1.1 | 45.9±3.8 | 40.6±0.9 | 53.6±2.1 | 37.3±2.1 | 42.0±3.9 | 44.9±2.2 | 48.9±7.4 | ** | 0.10 | 0.41 |
| $\text{WUE}_i$ | 57±6 | 70±9 | 45±5 | 91±12 | 46±9 | 73±5 | 87±22 | 109±23 | ** | * | 0.58 |
| $C_i/C_a$ | 0.72±0.03 | 0.72±0.04 | 0.77±0.02 | 0.64±0.04 | 0.80±0.03 | 0.69±0.02 | 0.63±0.07 | 0.55±0.07 | * | * | 0.48 |
| $C_c/C_a$ | 0.56±0.03 | 0.49±0.02 | 0.62±0.02 | 0.46±0.02 | 0.56±0.06 | 0.48±0.01 | 0.43±0.08 | 0.32±0.08 | ** | ** | 0.79 |

**Fig. 1.**
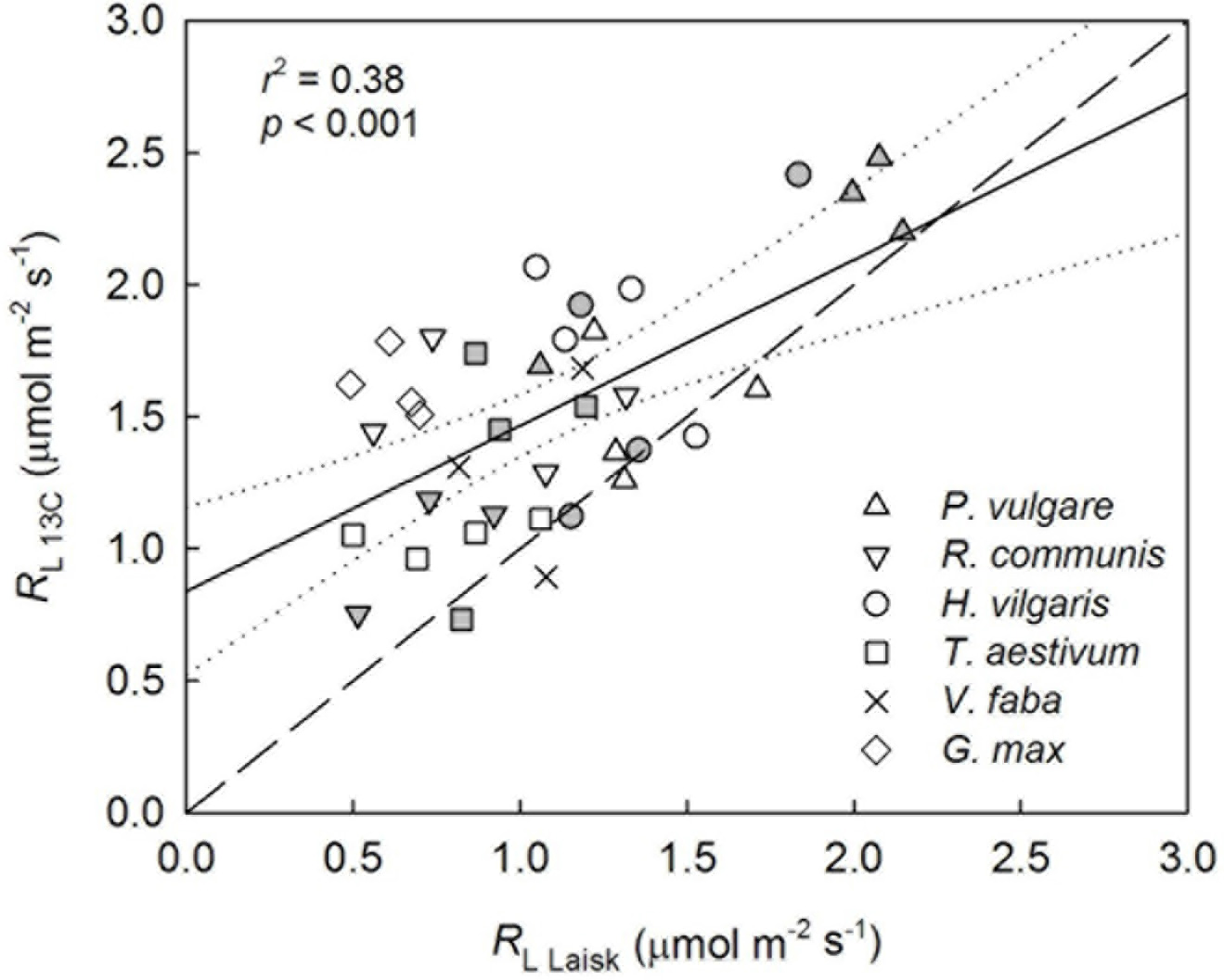
Correlation between respiration rate in light measured using the isotopic disequilibrium method (*R*_l 13C_) and the Laisk method (*R*_L Laisk_). Open symbols represent data of young leaves and filled symbols represent old leaves; *V. faba* and *G. max* were measured only on young leaves. Lines are the regression line (black solid line), upper and lower 95% confidence limits (dotted lines) and the 1:1 line (dashed line). Each symbol represents both parameter measured on the same leaf.

**Fig. 2.**
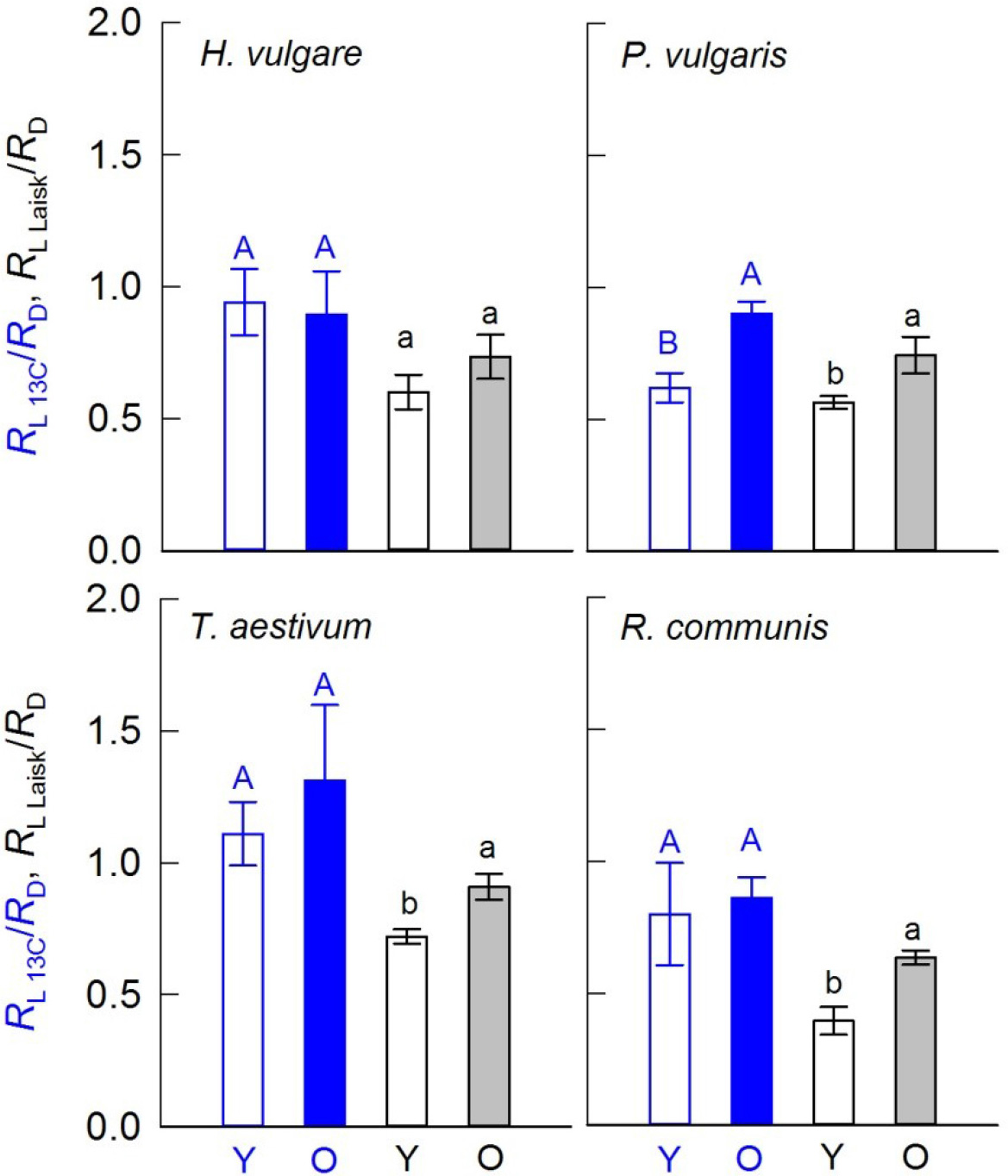
The ratio of respiration in light to that in darkness measured by the isotopic disequilibrium method (*R*_L 13C_/*R*_D_, blue bars) or the Laisk method (*R*_L Laisk_/*R*_D_, black bars) of young (Y, open bars) and old (O, filled bars) leaves. Different letters indicate significant difference between means within each species measured by the same method (*p*=0.05, *n*=4).

### Photosynthetic parameters

Leaf ageing had clear effects on many gas exchange parameters (Table 1) when averaged across species. Old leaves had an approx. 30% lower net CO_2_ assimilation rate (*A*), 58% lower stomatal conductance to water vapour (*g*_sw_), 47% lower mesophyll conductance (*g*_m_), 11% lower ratio of internal-to-atmospheric CO_2_ mole fraction (*C*_i_/*C*_a_), and a 19% lower ratio of chloroplastic-to-atmospheric CO_2_ mole fraction (*C*_c_/*C*_a_), as compared to young leaves. On the other hand, old leaves had a 13% higher 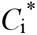 and 46% higher intrinsic water-use efficiency (WUE_i_ = *A*/*g*_sw_) compared with young leaves, averaged across species (Table 1). Nevertheless, *A* and *g*_m_ in *P. vulgaris* did not differ significantly between the two age classes. Across individual leaves of all species and age classes, *R*_L_ was not significantly correlated to *A*, *g*_sc_ or *g*_m_ (*r*^2^ <0.1, *p*>0.05), but was significantly correlated to *R*_D_, with both methods 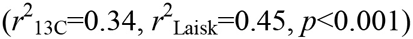.

### Isotope fractionation and mesophyll conductance

Carbon isotope discrimination during net CO_2_ assimilation (Δ_A_) showed clear differences during the two sets of online Δ measurements (Fig. 3), that is, the observed discrimination was influenced by the isotope composition of inlet CO_2_. This was due to the isotopic disequilibrium between respiratory (*R*_L_) and photosynthetic (*P*) CO_2_ fluxes. By contrast, Δ_p_ was not influenced by CO_2_ sources in any species (Fig. 3) supporting the accuracy of flux partitioning of *P* and *R*_L_.Furthermore, the calculation using Eqn 16 yielded estimates of *e* of −16.5‰ with ^13^C-depleted inlet CO_2_ and +11.2‰ with ^13^C-enriched inlet CO_2_ (averaged across species). Those estimates were close to values that could be simply computed from the δ^13^C difference between growth CO_2_ source and outlet CO_2_ (that is, *e* = δ_out_ − δ_growth CO2_ where δ_growth CO2_ = −10% and δ_out_ denotes the isotopic composition of CO_2_ in the leaf cuvette during measurements in light), assuming there was no fractionation between photosynthates and respired CO_2_ (Wingate *et al.*, 2007): *e* obtained in this way was −18.3% and + 6.1% with ^13^C-depleted and ^13^C-enriched inlet CO_2_, respectively. The agreement between the two calculations of *e* again indicates that our flux partitioning of *P* and *R*_L_ was performed properly.

**Fig. 3.**
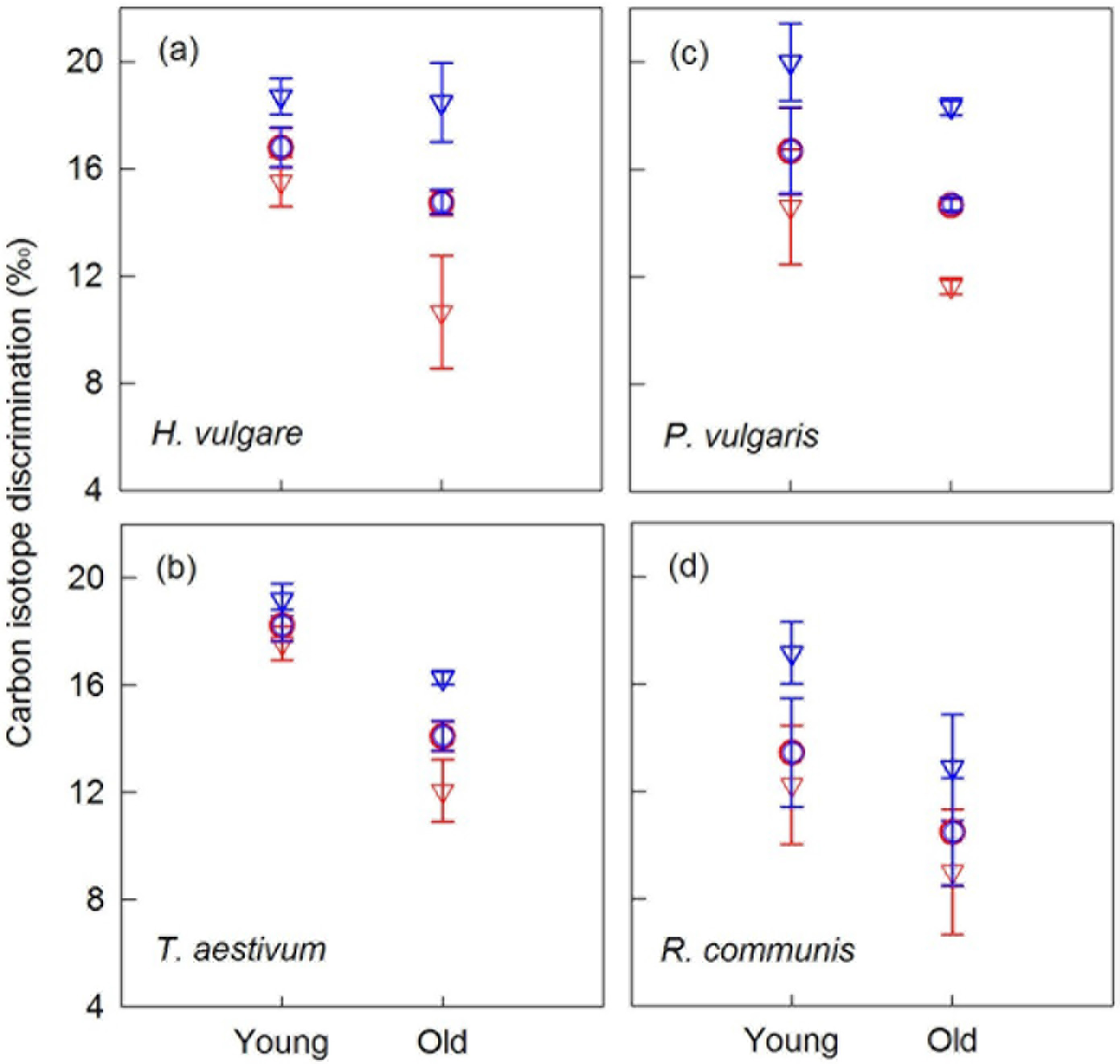
Carbon isotope discrimination during net CO_2_ assimilation (Δ_A_, triangles) and during apparent photosynthesis (Δ_P_, circles) measured using an ^13^C enriched CO_2_ source (red symbols) or a ^13^C depleted CO_2_ source (blue symbols). Panels a - d show data of individual species, and error bars are standard errors (*n*=4).

*g*_m_ was calculated from carbon isotope discrimination during apparent photosynthesis (Δ_p_) using equation (12). As measured under conditions similar to growth conditions using our isotopic disequilibrium method, *g*_m_ and *g*_sc_ showed a strong linear correlation across young and old leaves of all species (*g*_sc_=0.67*g*_m_+0.01, *r*^2^=0.82, *p*<0.001, Fig. S1). Meanwhile, *g*_sc_/*g*_m_ showed no significant correlation with *A* (*p*>0.05, *r*^2^ <0.1) or CO_2_ mole fraction in the leaf cuvette (*C*_out_, *p*>0.05, *r*^2^<0.1). The *g*_m_-*g*_sc_ relationship was used to calculate *g*_m_ of each leaf during Laisk measurements (*A*/*C*_i_ curves) and thus to calculate Γ* and *R*_L Laisk cc_ using *A*/*C*_c_ curves (cf. Fig. S4).This established that 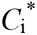 was generally lower than Γ* with a mean absolute difference of 5 μmol^-1^ for both young and old leaves (Fig. 4a), while *R*_L Laisk cc_ (obtained from *A*/*C*_c_ courves) was not different from *R*_L Laisk_ (obtained from *A*/*C*_i_ curves; Fig. 4b). An example of the offset in the common intersection point is given in Fig. S4.

**Fig. 4.**
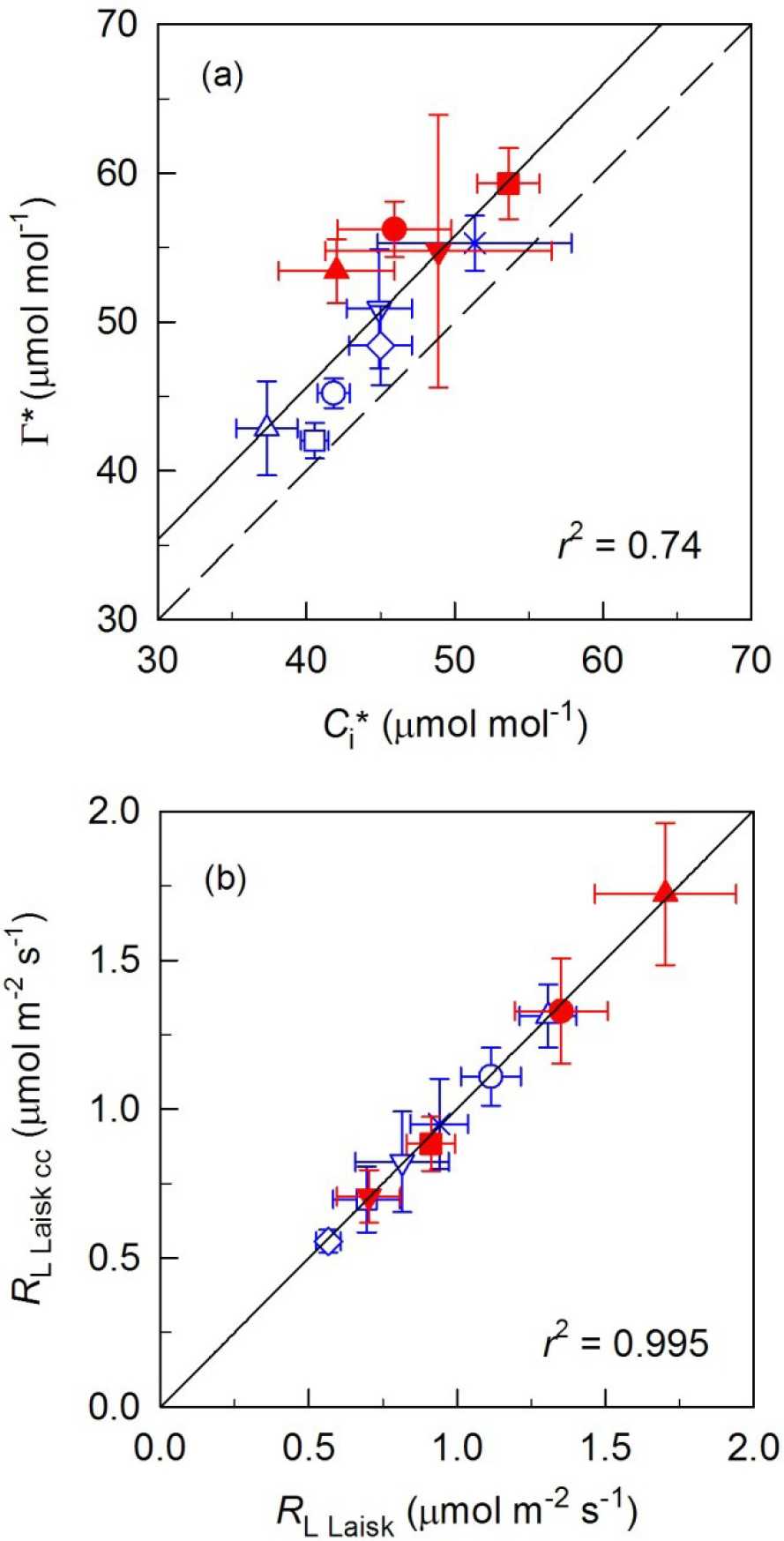
Relationship between (a) chloroplastic CO_2_ photocompensation point (Γ*) and apparent chloroplastic CO_2_ photocompensation point (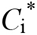), and between (b) respiration in light measured using modified Laisk method based on *C*_c_ (*R*_L Laisk cc_) and that measured using the Laisk method (*R*_L Laisk_). Species are separately marked with different symbols (see Fig. 1), blue symbols represent data of young leaves and red symbols represent old leaves. Black solid line is the regression line, and dashed line is the 1:1 line. Each symbol represents the mean and standard error of a species (*n*=4).

## DISCUSSION

In this work, *R*_L_ was measured using both an isotopic disequilibrium method and the classical Laisk method on single leaves of different species, and values obtained therefrom were compared.

### *Reliability of R*_L_ *values derived from isotopic disequilibrium*

The present results showed a positive correlation between the two sets of *R*_L_ measurements across all species and age classes, while on average R_L Laisk_ estimates were 28% smaller than *R*_L 13C_. To our knowledge, this is the first comparison of *R*_L_ estimated from the Laisk method and an isotopic disequilibrium method that does not require manipulation of photosynthetic gas exchange rates using non-physiological environmental conditions. It is not totally unexpected that the two methods provided consistently different *R*_L_ estimates, given that the measurements were performed with contrasting environmental conditions and different theoretical bases. The isotopic disequilibrium method measures CO_2_ efflux that is not labelled (i.e. respiration fuelled by old carbon) during leaf photosynthesis. An important assumption involved is that after a short period of labelling, no tracer (new carbon) has been incorporated into respiration. Any contribution of new carbon to the respiratory CO_2_ efflux will lead to an underestimation of *R*_L_. The potential error seems to be negligible, since our calculations using Eqn 8 (Table S2) showed that this assumption might have led to a 2.5% underestimation of R_L_ only, thus cannot explain the offset between *R*_L_ estimates measured by the two methods. Also, in perennial ryegrass, no new carbon was observed in shoot dark respiration for about 2 h following a 1 h-long labelling period (Lehmeier *et al.*, 2008), again suggesting insignificant underestimation of *R*_L_ by short-term labelling (30-45min). The labelling dynamics in shoot respiration should be similar to that of single leaves considering that leaf respiration contributes to about half of total plant respiration (Atkin *et al.*, 2007). However, information on labelling dynamics of single leaves is currently very limited, thus the kinetics of label appearance in day respired CO_2_ and its putative environmental dependence should be studied in a greater number of species.

### *Does R*_L Laisk_ *respond to environmental conditions imposed during measurement?*

Estimates of *R*_L_ differ between methods (Villar *et al.*, 1994; Yin *et al.*, 2011), and this effect is likely related to the different measurement conditions. Importantly, the response of *R*_L_ to environmental conditions like irradiance and CO_2_ concentration is not well understood to date, mainly due to methodological limitations. Light has long been recognized to inhibit *R*_L_ so that *R*_L_ is believed to be higher at very low light, a phenomenon that is possibly also at the origin of the Kok effect (Brooks & Farquhar, 1985; Villar *et al.*, 1994; Atkin *et al.*, 2000; Yin *et al.*, 2011). However, the effect of light at higher levels is not well documented. It is notable that both the Laisk and Kok method require manipulation of PAR, so the effect of PAR on *R*_L_ cannot be quantified with these methods. Also, uncertainty remains as to whether there is a short-term response of *R*_L_ to CO_2_ mole fraction. Early reports of a decrease of leaf *R*_D_ with short-term increase of CO_2_ (see the discussion by Amthor (2000) and Yin *et al.* (2011)), were suggested to be largely attributable to CO_2_ diffusive leaks during gas exchange measurements (Amthor, 2000; Jahnke & Krewitt, 2002; Long *et al.*, 2004; Gong *et al.*, 2015). Results of the short-term CO_2_ response of day respiration are scarce. However, using ^13^C-labelling, it was shown that respiratory metabolism (TCA pathway) increased as CO_2_ mole fraction decreased (Tcherkez *et al.*, 2008), while there seemed little effect on *R*_L_ assessed with the Kok method (Tcherkez *et al.*, 2012b). CO_2_ mole fraction can potentially impact on *R*_L_ via changes in nitrogen assimilation caused by altered rates of photorespiration (Tcherkez *et al.*, 2012a; Abadie *et al.*, 2016). On the one hand, increased photorespiration at low CO_2_ is believed to cause high mitochondrial NADH levels and thus inhibit TCA decarboxylases. On the other hand, the increased demand for carbon skeletons to assimilate nitrogen at high photorespiration should stimulate day respiratory metabolism (Abadie *et al.* 2016). However, the contribution of TCA decarboxylations to total respiratory CO_2_ production in the light is rather small when compared to pyruvate dehydrogenation (Tcherkez *et al.*, 2008). Therefore, the net effect of CO_2_ on *R*_L_ itself may be modest. Still, a short-term change in CO_2_ mole fraction may in principle influence *R*_L_, and thus the possibility that *R*_L_ is misestimated by the Laisk method cannot be excluded. This could contribute to explaining why Laisk estimates of *R*_L_ are smaller than ^13^C-derived estimates, as shown here.

Further, the low CO_2_ conditions used with the Laisk method may provoke a diffusive leak as the (non-controlled) CO_2_ concentration outside the cuvette is higher than inside. That would increase the estimate *R*_L_ if not accounted for properly, further affecting the relationship between *R*_L Laisk_ and *R*_L 13C_. In the present work, however, the leak effect was accounted for. Also, the leak coefficients of intact leaves (*K*_CO2_) measured here were generally very low, much lower than the producer-suggested value of 0.44 μmol s^-1^. Nevertheless, we found a clear leak artefact on *R*_L Laisk_ of *V. faba.* The diffusive leak had no significant effect on estimates of 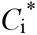 of young leaves of *V. faba* and *R. communis* (Fig. S2). Importantly, leak artefacts on *R*_D_ are also not ignorable given that measurements of *R*_D_ of small leaves are quite close to the detecting limit of currently available infra-red gas analysers. Since *K*_CO2_ may vary substantially between species and leave age classes, leak effects should be minimized (cf. Gong *et al.* 2017b) or accounted for by the measurement of the leak coefficient for every single leaf, as done here.

### Does g_m_ influence R_L_ estimates?

Another potential uncertainty associated with the *C*_i_-based Laisk method is the assumption on mesophyll conductance. The compensation point in the absence of day respiration, Γ*, is a *C*_c_-based value and thus should be determined from *A*/*C*_c_ curves rather than *A*/*C*_*i*_ curves. In other words, the use of *A*/*C*_i_ curves to estimate 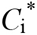 (as a proxy of *C*_c_) involves the assumption that *g*_m_ is infinite. Consequently, assuming an infinite *g*_m_ might lead to errors in the estimated Γ* and *R*_L_ by the Laisk method (von Caemmerer, 2013; Walker & Ort, 2015). Here, *g*_m_ of each leaf was quantified using online Δ measurements, and demonstrated that *g*_m_ of older leaves was 47% smaller than that of young leaves, in agreement with studies using both online Δ or florescence methods (reviewed in Flexas *et al.* (2008). The estimates of *g*_m_ obtained here were not very sensitive to errors in Γ*. In fact, the difference between species and age classes was not influenced by changes in Γ* within 20 μmol mol^-1^ (Fig. S3). Other methods like the constant J method were suggested to be sensitive to errors in Γ* (Harley *et al.*, 1992). Furthermore, the robust relationship between *g*_sw_ and *g*_m_ across all species found here was similar to that reported in tree leaves (Whitehead *et al.*, 2011). Knowing the relationship between *g*_sc_ and *g*_m_ allowed us to estimate *g*_m_ and thus convert *A*/*C*_i_ curves into *A*/*C*_c_ curves in the Laisk method. That way, we were able to derive the parameters of interest (Γ* and *R*_L_) from *A*/*C*_c_ curves (cf. Fig. S4). These calculations assumed that the *g*_sc_-*g*_m_ relationship was the same under the measurement condition of the Laisk method and normal growth condition, which is supported by the fact that *g*_sc_-to-*g*_m_ ratio showed no significant correlation with net assimilation rate or CO_2_ mole fraction in the leaf cuvette. Furthermore, analyses of published data also showed a strong *g*_sc_ − *g*_m_ relationship across species and growth conditions (Flexas *et al.*, 2013). Importantly, however, our results show that the Laisk method based on *A*/*C*_i_ curves systematically underestimated Γ* (by 5 μmol^-1^) but not *R*_L_ (i.e. *R*_L_ determined from *A*/*C*_i_ curves and *A*/*C*_c_ curves were identical).

Although statistical significance was found in *T. aestivum* only, the age effect on both 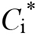 and the *C*_c_-based value of Γ* suggested that there was some error in the Laisk method. In fact, Γ* is given by [O_2_]/2*S*_*c*/*o*_ (where [O_2_] is oxygen mole fraction at carboxylation sites and *S*_c/o_ is Rubisco specificity) and is thus not expected to change with leaf age. Assuming a single conductance term from intercellular spaces (*C*_i_) to the site of carboxylation (*C*_c_) is perhaps not completely realistic, as some authors suggested that there is some resistance of the chloroplast envelope to intracellular CO_2_ movement (von Caemmerer, 2000), thereby leading to a lack of common intersection in Laisk curves (Tholen *et al.*, 2012). According to the model of Tholen *et al.* (2012), 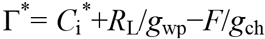, with total mesophyll conductance subdivided into conductance associated with cell wall and plasmalemma (*g*_wp_) and chloroplast envelope and stroma (*g*_ch_).Under such an assuption, the offset of 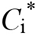 and apparent Γ* between age classes can be explained by a small increase in photorespiration *F* (according to the difference of *C*_c_) and a small decrease in *g*_ch_ with ageing. Improving the representation of mesophyll conductance in the Laisk method is beyond the scope of the present paper, but our results suggest that the estimates of Γ* or *R*_L_ obtained via the Laisk method are not precise enough (Gu & Sun, 2014), and should be viewed as approximations of actual Γ* and *R*_L_.

### Conclusions and perspectives

This study showed a high variation in *R*_L_ of similar leaves measured by both methods, and *R*_L_ was positively correlated to *R*_D_, but not to net CO_2_ assimilation rate or other parameters. These observations do not support the assumption that leaf *R*_L_ is a fixed proportion of photosynthesis or maximum *V*_c_ as used in many models (cf. De Kauwe *et al.* (2016)), but suggest that scaling *R*_L_ to *R*_D_ is a more reliable approach for the modelling purpose. We found a tendency for *R*_L_/*R*_D_ to increase during leaf aging, and this finding is not in agreement with that reported by Villar *et al.* (1995). The average age difference between young and old mature leaves was about 16-20 days in our study, much shorter than that of tree leaves (about 2 years) in the study of Villar *et al.* (1995). Taken as a whole, our results show that *R*_L_ estimates obtained using the isotopic disequilibrium method and the Laisk method were positively correlated, but *R*_L_ estimated by the isotopic disequilibrium method was generally higher than that measured by the Laisk method. Both methods captured the difference in *R*_L_ between species but found no effect of leaf ageing. Although *R*_L_ estimates differed between measurement techniques, most leaf-level studies (including the present study) support the notion that *R*_L_ is lower than *R*_D_ (Villar *et al.*, 1994; Yin *et al.*, 2011; Gong *et al.*, 2015; Tcherkez *et al.*, 2017a). Mesocosm-scale ^13^C labelling study also showed that stand *R*_L_ is inhibited by light (Schnyder *et al.*, 2003; Gong *et al.*, 2017a). Previous comparisons between Laisk and Kok methods showed a systematic difference between *R*_L_ estimates, with *R*_L_ estimated by the Kok method being generally lower than that measured by the Laisk method (Villar *et al.*, 1994; Yin *et al.*, 2011). Also in the case of the Kok method, it has been recently suggested that the apparent inhibition of respiration by light is at least partially explained by considerable changes in *C*_c_ during the manipulation of irradiance (Farquhar & Busch, 2017), in addition to other changes such as that in photochemical yield (for a review, see Tcherkez *et al.* 2017a b). Thus, our study suggests that common methods (Laisk or Kok) likely provide underestimated *R*_L_ values and thus overestimated inhibition of day respiration by light. For the mechanistic understanding of day respiratory metabolism, the response of *R*_L_ to light and CO_2_ mole fraction should be assessed in further studies, and the isotopic disequilibrium method is suitable for such a purpose since it does not require irradiance and CO_2_ alterations.

## ACKNOWLEDGMENTS

We thank The New Phytologist Trust for supporting the 18th New Phytologist Workshop ‘The Kok effect: beyond the artefact, emerging leaf mechanisms (KOALA)’ Angers, France, July 2016. We also thank all participants of this workshop for enlightening discussion and comments. This research was supported by the Deutsche Forschungsgemeinschaft (DFG SCHN 557/7-1).

### AUTHOR CONTRIBUTIONS

R.S. and X.Y.G. designed and planned the research; J.W. and R.S. performed the gas exchange measurements and isotope analyses; J.W., R.S. and X.Y.G. analyzed the data; G.T., R.S., H.S., and X.Y.G. discussed the results and implications; X.Y.G. wrote the first draft, and all authors contributed to the revision.

